# Incorporation of microbially salvaged urea-nitrogen into anabolic amino acids during hibernation in arctic ground squirrels

**DOI:** 10.1101/2025.10.10.681026

**Authors:** Sarah A. Rice, Kirsten Grond, Sarah M. Gering, Julie A. Reisz, Taylor R. M. Bailey-Parsons, C. Loren Buck, Khrystyne N. Duddleston

## Abstract

Hibernators rely on endogenous supplies of amino acids (AAs) and other nutrients to sustain organ function and skeletal muscle mass. Gut microbiota-mediated recycling of urea-nitrogen into AAs has long been considered a mechanism to assist in conserving nitrogen; however, the relevance of urea nitrogen salvage (UNS) to overall host physiology and metabolism is debated. We hypothesized that incorporation of microbially-liberated urea-nitrogen (MLUN) into AAs would be higher in host tissues of hibernating arctic ground squirrels compared to those of summer active squirrels, and that MLUN would support synthesis of anabolic AAs that regulate protein balance. To test this, we injected [^13^C, ^15^N_2_]-urea into summer squirrels and into squirrels hibernating at an ecologically relevant ambient temperature (−16°C). We found greater incorporation of MLUN into non-essential AAs and specific essential AAs in several tissues of hibernating squirrels compared to summer. We also observed increased ^15^N enrichment in leucine-isoleucine, citrulline and glutamine, anabolic AAs known to influence protein balance and trans-organ nitrogen balance. Compared to studies in which ground squirrels hibernated at ambient temperatures above 0°C, our results suggest that squirrels hibernating at subzero temperatures may up modulate synthesis of AAs that preserve protein and nitrogen balance during prolonged fasting and inactivity.

## Introduction

Hibernation is a highly regulated physiological, biochemical, and behavioral adaptation expressed across many species to promote survival during protracted periods of resource scarcity^1–4^. Obligate seasonal hibernators, such as ground squirrels, display short seasons of activity, when they reproduce and fatten in preparation for hibernation. Hibernation lasts 5 to 8 months depending on the species^3,5^, during which they neither eat nor drink and alternate between days or weeks of torpor and short periods of interbout arousal (IBA). In torpor, body temperature (T_b_) and metabolism are severely reduced, while during IBAs T_b_ and physiological processes return to euthermic levels, metabolic profiles change, and animals engage in gluconeogenesis and protein synthesis^2,4,6–8^. The arctic ground squirrel (*Urocitellus parryii*) is the most extreme obligate seasonal hibernator known. They hibernate for approximately eight months, and during bouts of torpor that can last over 21 days, they regulate core T_b_ to as low as −2.9°C, depress metabolism up to 98%, and emerge from hibernation with little muscle or bone loss^9–12^. The mean overwinter soil temperature of arctic ground squirrel hibernacula is −8.9°C, with minima as low as −25°C^12,13^. At these low ambient temperatures, arctic ground squirrels rely on both lipid and non-lipid (presumably amino acid) endogenous fuel sources^12^, and to date are the only obligate seasonal hibernator for which measures of Respiratory Exchange Ratio indicate use of mixed-fuel metabolism during hibernation^12^.

Urea, which is predominantly produced in the liver and released to systemic circulation, is formed from amino acid (AA) catabolism as a mechanism to excrete nitrogen waste from the mammalian body via the kidneys. Typically, between 1 and 80% of urea is excreted in urine depending on species and physiological state, with the remainder reabsorbed into the bloodstream^14,15^. Urea from mesenteric arterial blood flow crosses into all sections of the intestine where gut microbes expressing genes encoding for the urease enzyme (which is not present in the mammalian host) hydrolyzes urea to form free ammonia and carbon dioxide (Fig. 1A;^16^). Free ammonia is either taken up by the host or utilized by gut microbes as a substrate in microbial biosynthesis. Host re-use of microbially-liberated urea-nitrogen (MLUN), whether via uptake of free ammonia or of microbially derived N-containing metabolites, is referred to as urea nitrogen salvage (UNS). Microbial urease enzymes are widespread among microbial taxa^17^, and due to the high number of ureolytic microbes on the luminal surface of the cecum (an environment that likely selects for urease-positive microbes as a means to buffer pH), a strong concentration gradient forms between the blood and hindgut epithelium^16^; therefore, the cecum is considered to be the anatomical site most effective for urea nitrogen salvage in hindgut fermenters. Urea nitrogen salvage has long been studied in hibernating species^18–21^, and urea recycling has been documented in humans^22,23^ and other animals^16,24^.

**Fig. 1.**
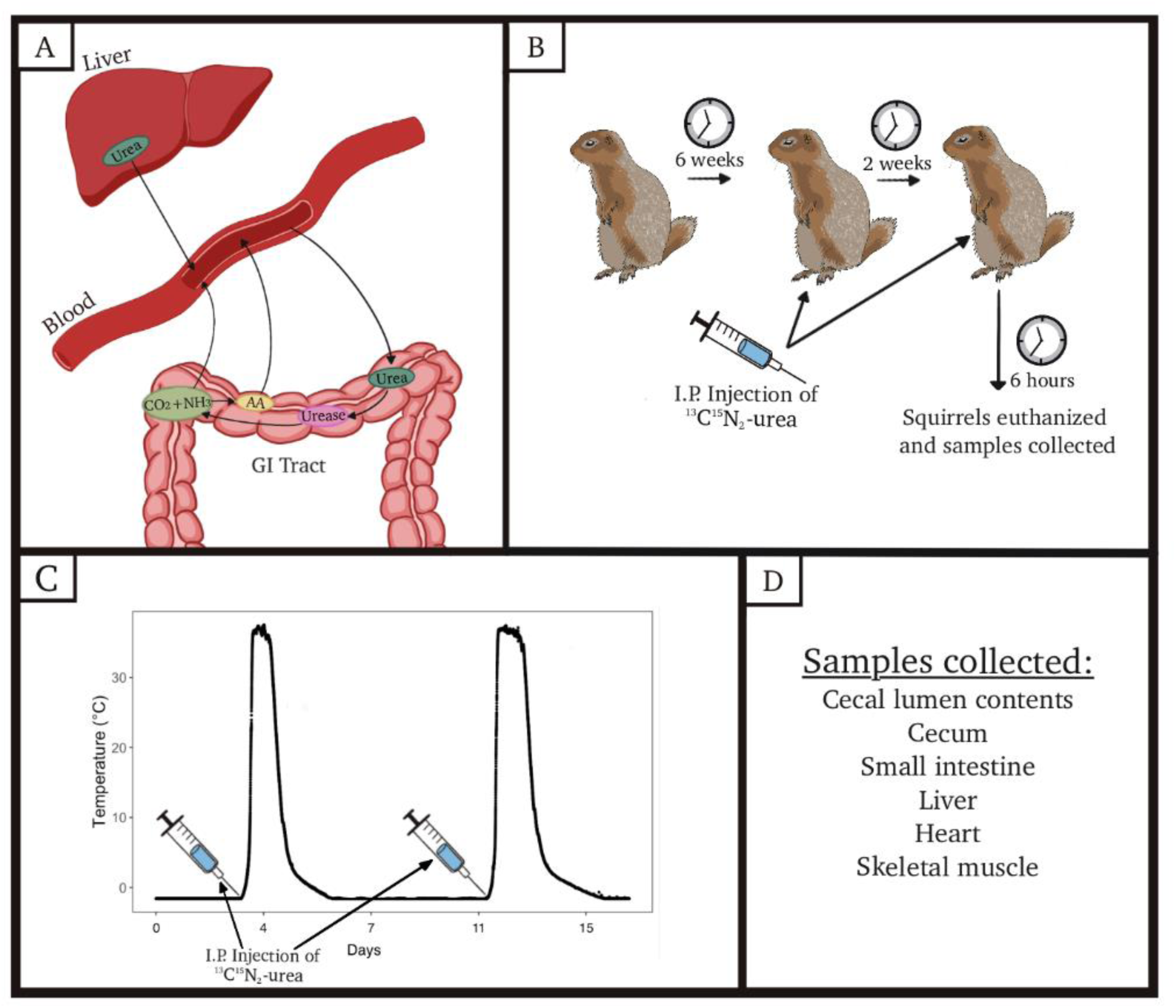
Experimental procedures. a. the proposed mechanism of urea nitrogen salvage. Gut microbes hydrolyze urea to form free ammonia and carbon dioxide. Ammonia can be utilized by the microbiota to form new amino acids, or be transported to the host via the portal blood stream. b. timing of [^13^C, ^15^N_2_]-urea injections in summer active arctic ground squirrels c. timing of [^13^C, ^15^N_2_]-urea during winter torpor bouts d. samples collected upon euthanasia

Given that arctic ground squirrels fast for up to eight months per year and rely on mixed-fuel metabolism during hibernation, their AA needs must be met through endogenous supplies, either via *de novo* synthesis or host-proteolysis. Many microbes can synthesize all AAs, including essential and non-essential AAs, and there is considerable interest as to the extent of host/microbiota exchange and the relevance of these AAs to host AA stores^25–27^. We have shown that UNS occurs in arctic ground squirrels ^21^, and Regan et al showed that 13-lined ground squirrels (*Ictidomys tridecemlineatus*) have increased capacity to recycle urea-nitrogen during hibernation and that metabolites from this process appear in the liver and skeletal muscle ^20^. Given this, a major question in hibernation physiology is the extent to which MLUN contributes to whole-body AA pools and protein sparing during long-term fasting.

Across mammalian species, whole body AA metabolism requires interdependence among organs, with some organs specializing in production of specific AA’s. For example, the small intestine is the major site of citrulline production and the kidney is the major site of *de novo* arginine production ^28–30^. Additionally, “anabolic” AAs, such as leucine, are known for regulating protein synthesis and/or breakdown, insulin sensitivity, and nitrogen balance^30^. We have shown previously that multiple organs appear essential for incorporating free nitrogen into non-essential AAs (NEAAs) and essential AAs (EAAs; including the branched-chain EAA leucine) during hibernation via non-microbial metabolic pathways^31^. Additionally, Chanon et al. (2018) demonstrated that serum from hibernating bears exerts a protein sparing effect on cultured muscle protein^32^. Protein sparing is potentially more important in hibernators experiencing extreme temperatures that induce protein catabolism^12,33^; however, the mechanism is not fully understood and circulating anabolic AAs are an intriguing potential contributor to this process^32^.

We aimed to investigate whether anabolic AA metabolism pathways utilizing MLUN were upregulated across organs in hibernation as compared to during the active season. We were specifically interested in the potential interdependence of MLUN and inter-organ dynamics. During hibernation gene expression across tissues is dramatically altered^34,35^, AA stores are retained^31,36^, and UNS occurs^20,21,37^; therefore, increased inter-organ utilization of MLUN during hibernation could represent a synergy between microbial adaptation in hibernation and innate host mechanisms to increase AA synthesis in organs. We hypothesized that incorporation of MLUN into EAAs and NEAAs would be higher in tissues of hibernating arctic ground squirrels compared to those of summer active squirrels. We also hypothesized that MLUN would support synthesis of anabolic AAs through UNS. In contrast to the study by Regan et al, we aimed to understand the potential use of MLUN in additional organ systems and across a broader metabolome, and in particular at temperatures that are more ecologically relevant to arctic ground squirrels than what has been previously published^20^.

## Methods

### Animal Capture

Free-living juvenile arctic ground squirrels (∼50:50 M:F) were live-trapped (Tomahawk [Hazelhurst, WI] baited with carrot) during the summers (early July) of 2017 and 2019 on the northern side of the Brooks Range in Alaska near the Atigun River (68°38′N, 149°38′W). Squirrels were transported to the University of Alaska Anchorage within 1 week of trapping and quarantined for 2 weeks.

### Animal Husbandry

Following quarantine, squirrels trapped in 2019 (used for hibernation experiments in winter 2020-2021) were surgically implanted with temperature-sensitive transmitters (TA10TA-F40; Data Sciences International [DSI], Holliston MA) to enable continuous monitoring of hibernation in real time^38,39^. Squirrels trapped in 2017 (used for summer experiments in 2018) were not implanted. All squirrels were housed individually in 18 ¾” W x 12 ¼” L x 12 ¼” D stainless steel wire mesh hanging cages hung over ammonia-absorbing litter, with cotton nesting material and metal tubes for enrichment. Squirrels were housed at ambient temperature (T_a_) of 18-19°C and 12L:12D hr light/dark cycle, and fed a standard rodent chow diet (Mazuri, St. Louis, MO, USA) with the addition of ∼5g of apples daily, and water *ad libitum* until the start of hibernation. In addition, squirrels trapped in 2017 were administered prophylactic metronidazole upon transfer to the UAA vivarium (mid July 2017; 0.5g in 25mL sweetened water/day for 10 days) to prevent overgrowth of *Clostridium difficil*e, followed by a 4 - 8 week recovery period after which they hibernated over the 2017-2018 winter (see below) prior to use in experiments in summer of 2018. Although metronidazole alters the gut microbiota, in particular affecting members of the family *Lachnospiraceae*, the community returns to a composition similar to pre-treatment within 2 - 4 weeks^40^. Analysis of the cecal microbiota of the 2017 squirrels at the time of euthanasia (in 2018; see below) revealed a community composition typical of those reported previously^41^ ^42,43^) with multiple members of the *Lachnospiraceae* in the top 25 most abundant genera (Supplementary Fig. 1; e.g. *Blautia, Coprococcus, Dorea, Roseburia, Ruminococcus*). Prophylactic metronidazole was not indicated for squirrels trapped in 2019. Squirrels trapped in 2019 were allowed to hibernate over the 2019-2020 winter and used in experiments in 2020-2021.

At the end of the first active season in captivity when squirrels showed evidence that they were ready to hibernate (decrease in activity and food consumption; curled hibernation posture), they were transferred to Nalgene™ tubs (15” W x 22” L x 8” D) with wood shavings, cotton nest material and steel wire lids, and held in the dark in an environmental chamber at T_a_ = 2°C with neither food nor water. During their first hibernation season in captivity (which occurred prior to the onset of experiments), squirrels trapped in 2017 and 2019 were checked daily for signs of arousal using the sawdust method^44^. Squirrels trapped in 2019 were monitored electronically for hibernation stage via DSI temperature transmitters during their second hibernation season^44^. All animal handling and research was conducted under approved IACUC protocols (769548, 842836, 1353732, 1428231).

### Experimental Design

Following one hibernation season in captivity, yearling squirrels were transferred to hanging wire cages and held under active euthermic conditions as described above. Squirrels were fed experimental diets as described below.

#### Summer active squirrels

Squirrels trapped in 2017 and overwintered in 2017-2018 were assigned to two diet treatment groups upon exiting hibernation in 2018: those fed 9% dietary protein content (n = 8 males and 7 females; Teklad diet TD.170165; Envigo [now Inotiv], Madison, WI, USA) and those fed 18% dietary protein (n = 9 males and 13 females; Teklad diet TD.170164). These diet manipulations took place between April and July, 2018. To increase the palatability of the diets, prevent caching and loss of food through the wire bottoms, and ensure squirrels consumed all food, food pellets (∼50g/day) were blended with Karo syrup (<1%) and just enough water to create a mush. The addition of Karo syrup resulted in only minor changes to the calorie and macronutrient content of the diets (Supplementary Table 1). After six weeks on their respective diets, summer active squirrels were injected intraperitoneally (I.P.) with [^13^C, ^15^N_2_]-urea (< 1ml; 50 mg/kg; ^13^C, 99%; ^15^N/^15^N, 98%; Cambridge Isotope Laboratories, Inc; Fig. 1C). Two weeks later (after 8 weeks on their respective diets) a second injection was given. Approximately 6.5-8.5hrs following the second urea injection (during which no food was given), squirrels were euthanized by overdose of isoflurane (5% mixed with medical grade oxygen and delivered at 1.5L/min) followed by decapitation, and tissues collected as described below.

Despite prophylactic administration of metronidazole to juveniles trapped in 2017 before hibernation, four yearlings developed gastrointestinal dysbiosis (*Clostridium difficile)* during the study. We injected these squirrels subcutaneously with saline solution (0.9% NaCl, Butler Schein) and syringe fed them their experimental diet until they recovered. Once recovered, the squirrels were “reset” to experimental time-zero, resuming the diet manipulation for 6 weeks until their first urea injection.

#### Hibernating squirrels

Squirrels trapped in 2019 and overwintered in 2019-2020 were divided into two dietary treatment groups upon exiting hibernation in 2020: either 9% (5 males + 7 females; Teklad diet TD.170165) or 18% (4 males + 4 females; Teklad diet TD.170164) protein ^45^. Squirrels were fed ∼45g of pellets daily, which was converted into mush by adding water and 1 ml (0.92 g) of sunflower oil to prevent caching and ensure all food was consumed. We used sunflower oil rather than Karo syrup as it better improved diet palatability, the addition of which resulted in minor shifts in the calorie and macronutrient content of the diet (Supplementary Table 1). Squirrels were also given ∼5g of apples daily and water *ad libitum*. Squirrels were fed these diets for the remainder of the 2020 active season until they re-entered hibernation and were housed as described previously except at T_a_ = −16°C (T_a_ known to induce reliance on mixed-fuel metabolism in arctic ground squirrels ^12,45^).

After at least 90 days of cumulative torpor (i.e., days in torpor exclusive of IBAs; between January and February, 2021) and within 2 days of the next predicted natural IBA, torpid squirrels were transferred to a warm room (19-20°C), immediately given an I.P. injection of [^13^C, ^15^N_2_]-urea (n = 50 mg/kg) and then artificially aroused to eliminate the introduction of variation in the event that injection caused some but not all squirrels to arouse (Fig. 1B). The timing of IBAs is presented in Grond et al (2023), and was predicted by determining the average torpor bout length for each squirrel, which is consistent across the majority of hibernation ^12,39,45^. Squirrels were artificially aroused by placing them in a warm room (19-20°C) until they displayed evidence that they were no longer in torpor (i.e. loss of hibernation posture and movement in their hibernation tub; ∼5h). When first transferred to the warm room we gently manipulated their limbs and rubbed their abdomens for 0.5 to 1hr. Once squirrels displayed evidence they were no longer in torpor they were returned to the hibernation chamber (−16°C) to reach euthermic temperatures (> 35°C), complete the IBA, and subsequently passively cool to re-enter torpor. A second injection of labeled urea was administered within 2 days of the next predicted IBA, and squirrels were artificially aroused and then returned to the hibernation chamber following the protocol of the first injection. The vast majority of metabolic activity occurs when animals are at euthermic temperatures^46^; therefore, for statistical comparisons between summer active and hibernating squirrels, we only included data from hibernating animals that were at euthermic temperatures (at or above 35°C) for 6.75 to 9 hr during the IBA that followed their second urea injection. This reduced our overall sample size for hibernating animals to n=7 (see the statistical section below), but ensured both summer active and hibernation groups were euthermic for similar amounts of time post-second injection. After the squirrels had returned to torpor (after the second injection), they were euthanized by decapitation and sampled for tissue and cecal contents as described below and shown in Fig. 1D.

### Sample collection

Samples from summer active squirrels were collected immediately following euthanasia and held on ice in cryotubes until transferred to −80°C within 1 hr. Samples collected from hibernating squirrels were snap-frozen in liquid N_2_ and held on ice prior to transfer to −80°C within 1 hr. To collect contents from the lumen of the cecum, the cecum was removed and lumen material gently squeezed into a sterile petri dish prior to transfer to a cryotube. Subsequently, the cecum was opened and the remaining lumen material was rinsed off using sterile saline solution. The mucosal layer was removed by scraping the inside of the cecum with a glass slide, and tissues of the cecum (epithelium plus muscle) were collected. Samples of the small intestine (ileum), liver (left lobe), skeletal muscle (left quadricep) and heart (apex tip) tissues were collected. All tissues were thoroughly rinsed with ice-cold sterile saline until they ran clear prior to placement in cryotubes.

### Ultra-High-Pressure Liquid Chromatography-Mass Spectrometry (UHPLC-MS) Metabolomics

Metabolomics and fluxomic analysis were performed as previously described^11,31,47,48^. Briefly, all samples were extracted using ice cold solution of methanol:acetonitrile:water (5:3:2) at a concentration of 15 mg/mL. Samples were agitated with a bead beater for 5 min, vortexed vigorously for 25 min, and then centrifuged at 18,000 g for 15 min, all performed at 4°C. Supernatant extracts (10 µl) were analyzed on a Thermo Vanquish UHPLC coupled to a Thermo Q Exactive mass spectrometer (MS) using a 5 min C18 gradient and electrospray ionization as previously described ^49^. Data were acquired across a scan range of 65-975 m/z with 4 kV spray voltage, 15 sheath gas and 5 auxiliary gas; separate positive and negative ion polarities were run as described previously ^47,48^. During analysis, blanks and QCs (which are pooled from all samples) were interspersed throughout the run between samples to correct for drift and background. Peak picking and metabolite assignment were performed using Maven (Princeton University) alongside the KEGG database ^50,51^, confirmed against chemical formula determination from isotopic patterns and accurate mass, and validated against experimental retention times for >650 standard compounds from our in-house metabolite library (Sigma Aldrich; MLSMS, IROATech, Bolton, MA, USA). Leucine and isoleucine are reported jointly, as a co-eluted signal.

### Data Analysis

Data were analyzed with MetaboAnalyst 5.0 (www.metaboanalyst.ca) ^52^ and graphed using GraphPad Prism (v. 10, GraphPad Software). Data show the percentage of ^15^N enrichment from isotopically labeled urea into metabolites, original data means, and 95% confidence intervals. Percentage enrichment is calculated by ^15^N metabolite peak area/(^15^N metabolite peak area + ^14^N metabolite peak area) x 100 following natural abundance correction of ^15^N (0.364%). This type of normalization allows comparison between different metabolite batch runs. Data were also square root transformed and autoscaled for normalization within MetaboAnalyst. Normalizations were visually inspected within the software, square root transformation was chosen as the method of transformation based on these visualizations. For metabolites with more than one nitrogen atom, only one ^15^N nitrogen is reported. Since specimens from hibernating and summer active squirrels were analyzed in separate batches, we compared ^15^N incorporation into only the metabolites that were observed in both batches (Supplemental Data file 1).

Data were statistically analyzed using ANCOVA. As noted above, squirrels were fed diets of 9% or 18% protein; however, because the overarching goal of the study reported here was to compare hibernating squirrels to summer active squirrels, we used diet as a covariate factor to control for dietary differences in all our analyses. While the influence of diet on fluxomics and metabolomics in arctic ground squirrels was beyond the scope of this analysis, we previously examined the effect of diet on gene expression during hibernation and found no effect on the expression of urease genes ^45^. We found no influence of sex on ^15^N enrichment and thus did not include sex as a covariate in our analysis.

^15^N Metabolite profiles of tissue samples were visualized using non-metric multidimensional scaling (NMDS) based on Bray–Curtis dissimilarities, using the vegan R package (2.7.1). To assess the influence of dietary protein and tissue type on ^15^N metabolite profiles, permutational multivariate analysis of variance (PERMANOVA) was conducted using the adonis2 function in the vegan R package.

## Results

### Isotopically labeled nitrogen derived from urea is integrated into amino acids in the cecum lumen contents of arctic ground squirrels

Given that the cecum lumen contains the highest densities of microbes in the gastrointestinal tract of hindgut fermenters ^16^, we analyzed cecum lumen content from hibernating and summer active arctic ground squirrels for amino acids (AAs) and other metabolites containing ^15^N derived from isotopically-labeled urea. We found ^15^N enrichment in EAAs, NEAAs and metabolites such as nucleobases and 5-oxoproline in the cecum lumen content of both hibernating and summer active squirrels (Fig.2). Greater ^15^N enrichment was observed in the NEAAs glutamate (p<0.0001), and aspartate (p=0.0058) (Fig. 2a), the essential branched-chain amino acids (EBCAA) valine (p=0.0118, Fig. 2b), and the nucleobase adenine (p<0.0001) (Fig. 2c) of hibernating squirrels as compared to summer active squirrels. While not significantly different, there was modest incorporation of ^15^N into the EAAs methionine, lysine, tryptophan and phenylalanine in both active and hibernating squirrels (Fig. 2b). Additionally, ^15^N-labeled glutamate, aspartate, 5-oxoproline and adenine typically accounted for well over 1% of their total metabolite concentrations in the lumen content of hibernating squirrels (Fig. 2).

**Fig. 2.**
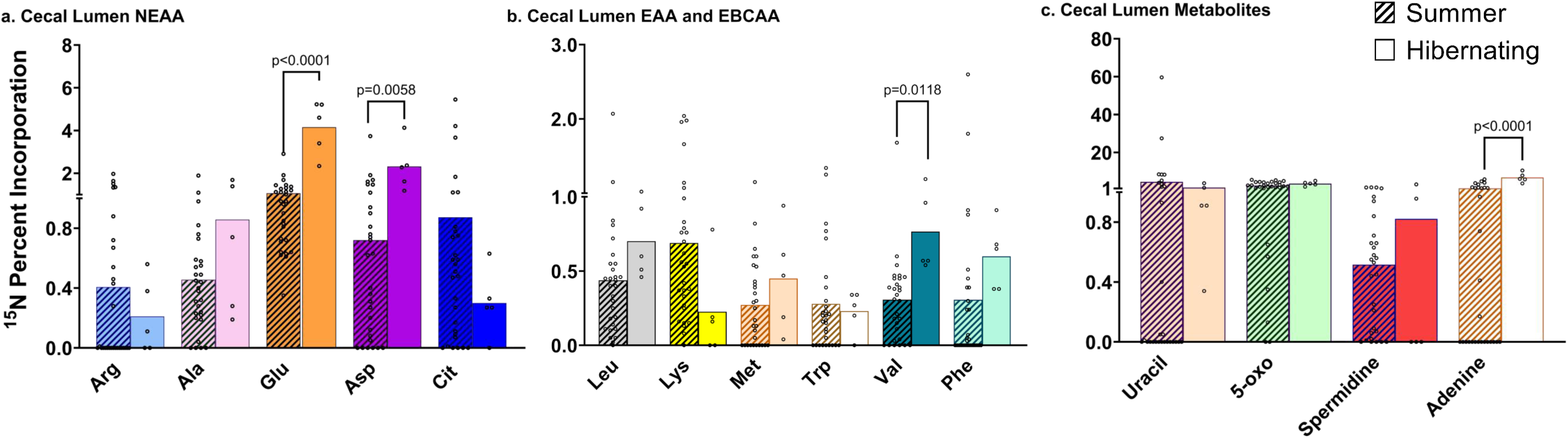
Urea-derived ^15^N percent enrichment into NEAAs and EAAs was higher in the cecal lumen of hibernating arctic ground squirrels. Enrichment of ^15^N in glutamate, aspartate, valine and adenine was higher in the lumen contents of hibernating squirrels, whereas incorporation into lysine trended higher in that of summer-active squirrels. a.) NEAA ^15^N enrichment. b.) EAA and EBCAA ^15^N enrichment. c.) metabolite ^15^N enrichment. ^15^N enrichment is presented as a percentage of the total metabolite, data show mean 95% confidence interval. Hatched bars indicate summer squirrels (n = 29), solid bars indicate hibernating squirrels (n= 5). Statistical analyses conducted only for metabolites detected in both hibernation and summer samples via multivariate linear regression, with diet as a covariate (p<0.05).

### More ^15^N-labeled metabolites were incorporated into GI tract tissues from hibernating squirrels compared to summer active squirrels

We found ^15^N enrichment into NEAA, EAA and other metabolites in cecal tissue of both hibernating and summer active squirrels, though there was greater enrichment in the NEAA glutamine (p<0.0001) in cecum tissue from hibernating squirrels compared to summer active squirrels (Fig. 3a). Additionally, ^15^N incorporation in the purine nucleotide UMP was higher in cecal tissue from hibernating compared to summer active squirrels (p<0.0001, Fig. 3c). While not significantly different between physiological states, ^15^N in citrulline was especially high in cecal tissue, reaching over 10% in many animals (Fig. 3a, c). Additionally, ^15^N in the EAAs methionine and threonine was present in cecal tissue of both hibernating and summer active squirrels, albeit below 1% (Fig. 3b).

**Fig. 3.**
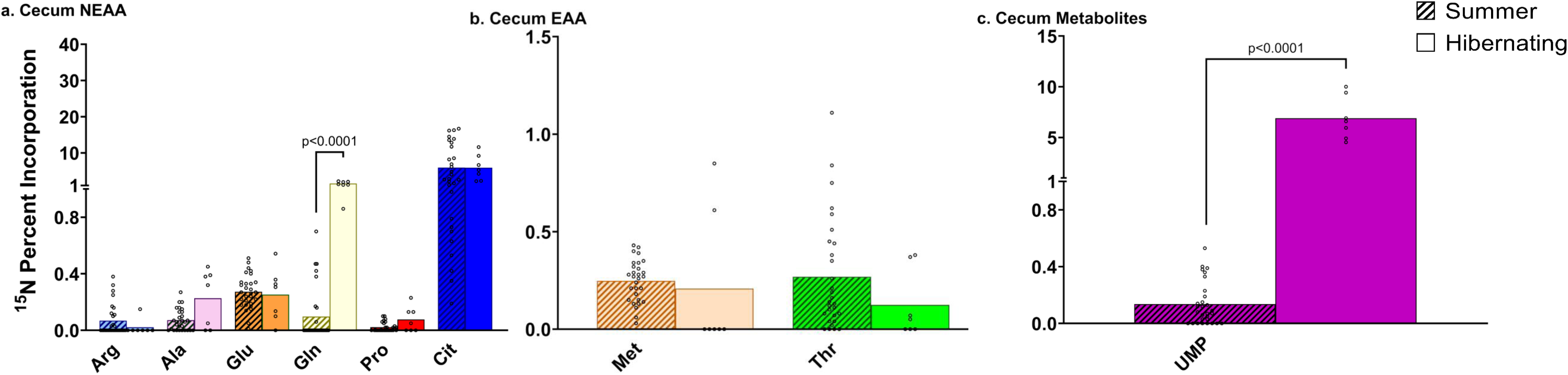
Urea-derived ^15^N enrichment in NEAA was higher in the cecum tissue of hibernating arctic ground squirrels. Enrichment of ^15^N into glutamine, and UMP was higher in the cecum tissue of hibernating squirrels a.) NEAA ^15^N enrichment. b.) EAA and EBCAA ^15^N enrichment. c.) metabolite ^15^N enrichment. ^15^N enrichment is presented as a percentage of the total metabolite, data show mean 95% confidence interval. Hatched bars indicate summer squirrels (n = 29), solid bars indicate hibernating squirrels (n= 6). Statistical analyses conducted only for metabolites detected in both hibernation and summer samples via multivariate linear regression, with diet as a covariate (p<0.05).

The small intestine is a major site for AA utilization and transport to the periphery, as well as for citrulline production in many mammalian species ^53^; therefore, we analyzed this tissue for the presence of ^15^N-labeled AA and metabolites. We found ^15^N metabolite enrichment in the small intestine in both summer active and hibernating squirrels, but with greater amounts observed for most metabolites in hibernating squirrels. Alanine (p=0.00182), citrulline (p=0.0182) and leucine-isoleucine (p<0.0001) were higher in the small intestine of hibernating squirrels as compared to summer active squirrels (Fig. 4). Conversely, glutamate was higher in summer squirrels compared to hibernating squirrels (p=0.0011, Fig 4a). Lastly, leucine-isoleucine overall showed the greatest amount of ^15^N incorporation in the small intestine (Fig. 4b).

**Fig. 4.**
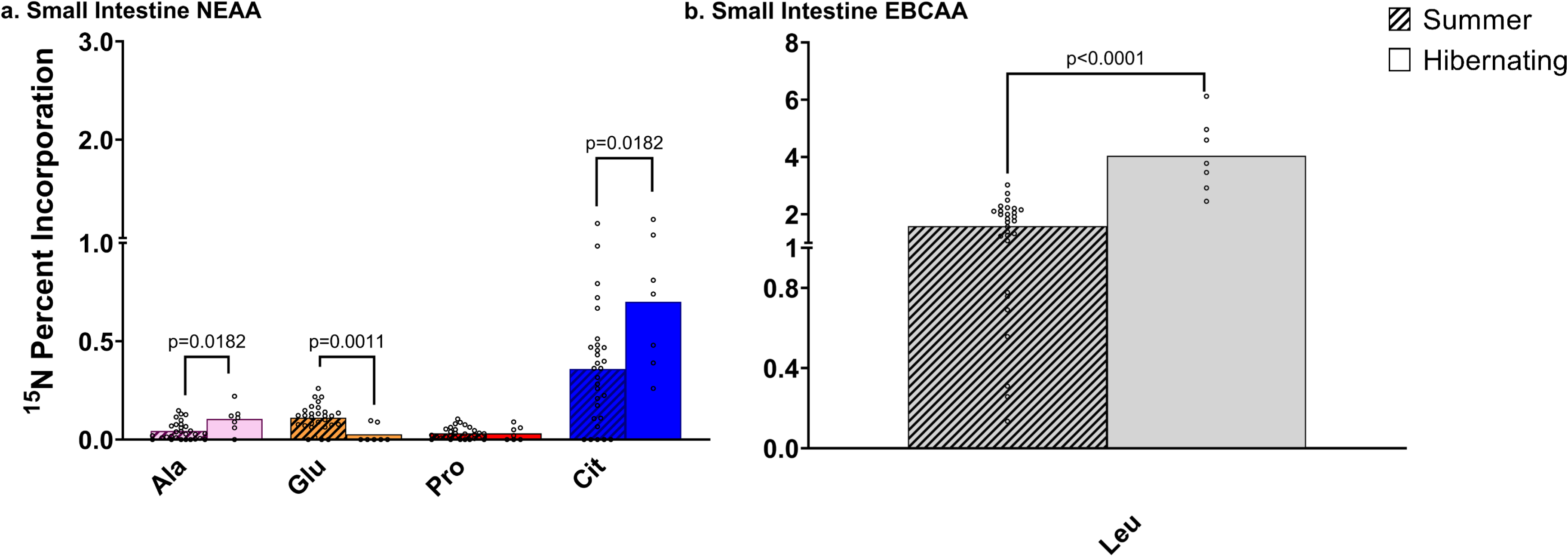
Urea-derived ^15^N enrichment in NEAA and EBCAA in the small intestine of arctic ground squirrels. Enrichment of ^15^N into alanine, citrulline and leucine-isoleucine was higher in hibernating squirrels, whereas glutamate was higher in summer squirrels. a.) NEAA ^15^N enrichment. b.) EBCAA ^15^N enrichment. ^15^N enrichment is presented as a percentage of the total metabolite, data show mean 95% confidence interval. Hatched bars indicate summer squirrels (n = 29), solid bars indicate hibernating squirrels (n= 7). Statistical analyses conducted only for metabolites detected in both hibernation and summer samples via multivariate linear regression, with diet as a covariate (p<0.05).

### More ^15^N enrichment in EAAs and NEAAs occurred in peripheral tissues of hibernating squirrels compared to summer active squirrels

Metabolites and ammonia derived from gut microbes and gut tissue are transported to the liver via the portal vein, and as such, any influence or appearance of ^15^N in host peripheral systemic/organ level metabolism may be expected to take place first in the liver. We found greater ^15^N enrichment in alanine (p<0.0002), glutamate (p<0.0001), glutamine (p<0.0001), glycine (p=0.0294), leucine-isoleucine (p<0.0001), and the deoxynucleoside thymidine (p<0.0001), in the liver of hibernating vs. summer active arctic ground squirrels (Fig. 5). Conversely, there was greater ^15^N-enriched methionine (p=0.0006) and threonine (p=0.006) in the liver of summer active vs. hibernating squirrels (Fig. 5b). Overall, leucine-isoleucine, glutamine, and thymidine all show the highest levels of ^15^N incorporation in hepatic tissue compared to other labeled metabolites (Fig. 5).

**Fig. 5.**
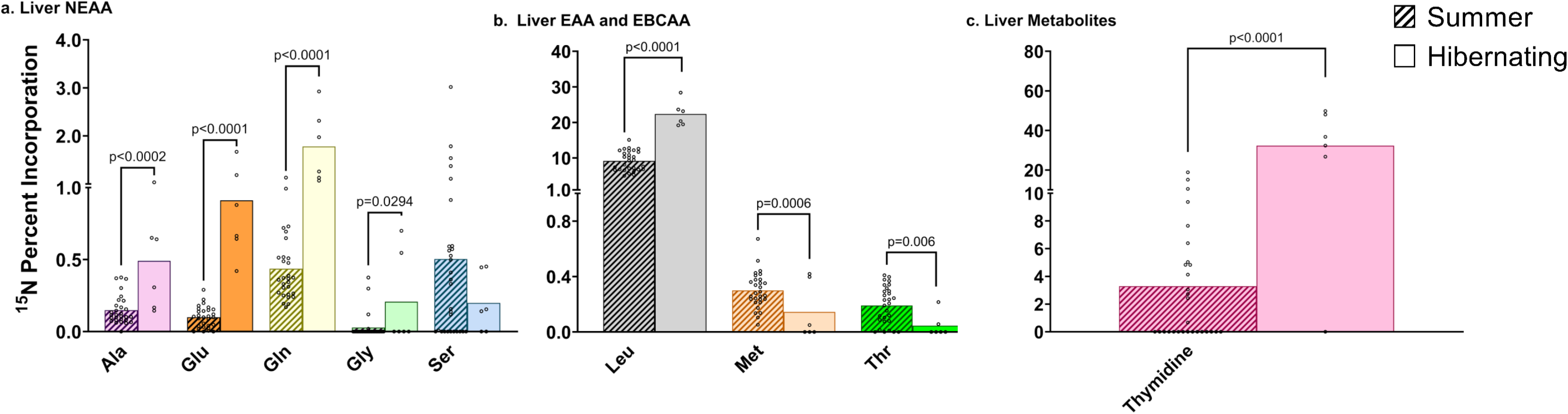
Urea-derived ^15^N enrichment in NEAA and EBCAA was higher in the liver of hibernating arctic ground squirrels. Incorporation of ^15^N into alanine, glutamate, glutamine, glycine, leucine-isoleucine, and pyrimidine deoxynucleoside thymidine was higher in hibernating squirrels, whereas enrichment in methionine and threonine was higher in summer squirrels. a.) NEAA ^15^N enrichment. b.) EAA and EBCAA ^15^N enrichment. c.) metabolite ^15^N enrichment. ^15^N enrichment is presented as a percentage of the total metabolite, data show mean 95% confidence interval. Hatched bars indicate summer squirrels (n = 29), solid bars indicate hibernating squirrels (n= 6). Statistical analyses conducted only for metabolites detected in both hibernation and summer samples via multivariate linear regression, with diet as a covariate (p<0.05).

Because blood leaving the liver travels first to the heart via the hepatic vein and inferior vena cava, and then is distributed throughout the body to peripheral skeletal muscle, we examined heart and skeletal muscle for incorporation of ^15^N into tissue metabolites. We found higher levels of ^15^N in glutamine (p=0.0110) and leucine-isoleucine (p<0.0001) in the heart tissue from hibernating arctic ground squirrels compared to summer active squirrels. While not differing between physiological states, ^15^N -arginine, -methionine, and -alanine were also observed in heart tissue, and leucine-isoleucine had the highest percentage of ^15^N enrichment (Fig. 6). In skeletal muscle, which has slow protein synthesis rates ^54^ and is one of the most distant tissues from the GI tract, there were higher amounts of ^15^N incorporated in glutamine (p<0.0001), and citrulline (p=0.0090) in hibernating squirrels as compared to summer active squirrels (Fig. 7).

**Fig. 6.**
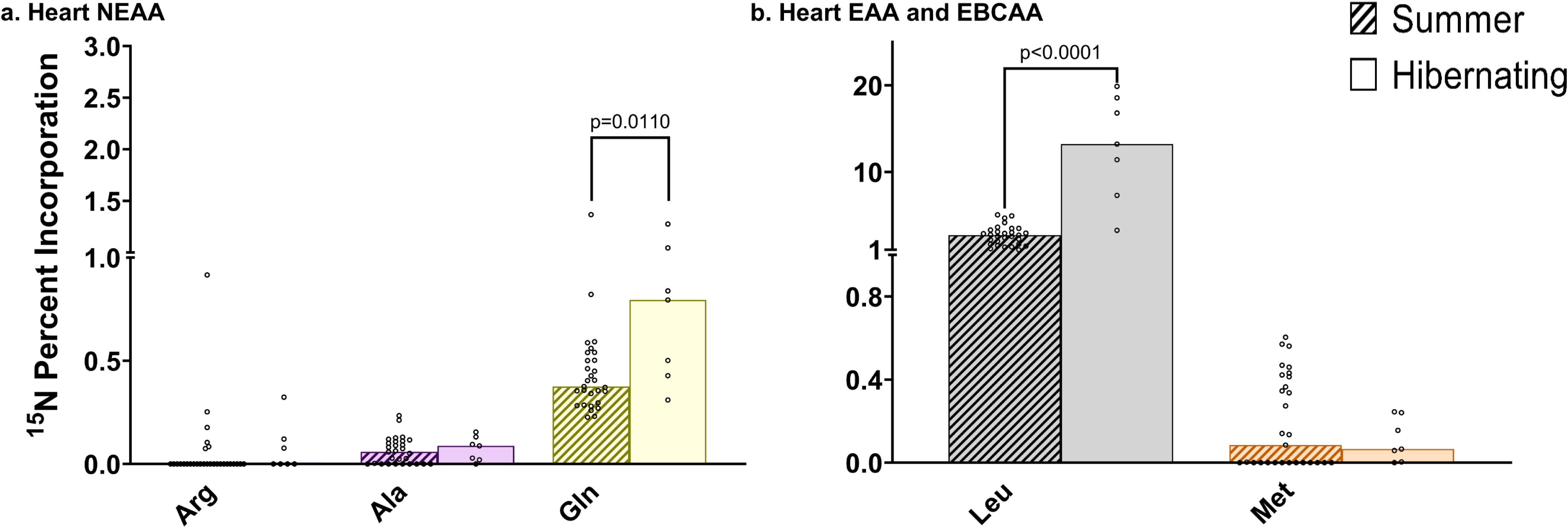
Urea-derived ^15^N enrichment in NEAA and EBCAA was higher in the heart of hibernating arctic ground squirrels. Enrichment of ^15^N into glutamine and leucine-isoleucine was higher in the heart of hibernating squirrels. a.) NEAA ^15^N enrichment. b.) EAA and EBCAA ^15^N enrichment. ^15^N enrichment is presented as a percentage of the total metabolite, data show mean 95% confidence interval. Hatched bars indicate summer squirrels (n = 29), solid bars indicate hibernating squirrels (n= 7). Statistical analyses conducted only for metabolites detected in both hibernation and summer samples via multivariate linear regression, with diet as a covariate (p<0.05).

**Fig. 7.**
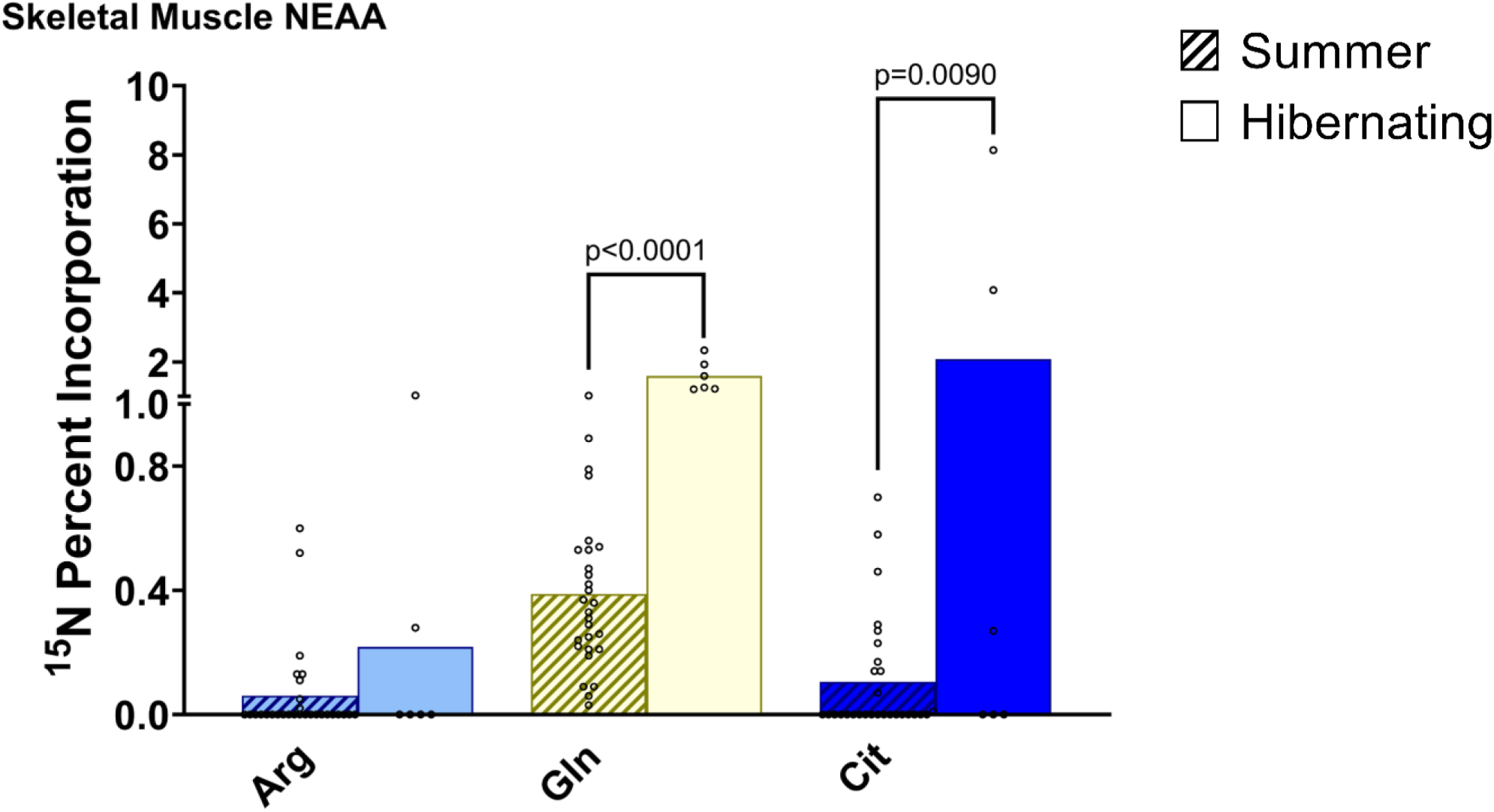
Urea-derived ^15^N enrichment into NEAA was higher in the skeletal muscle of hibernating arctic ground squirrels. Enrichment of ^15^N into glutamine and citrulline was higher in the skeletal muscle of hibernating squirrels. ^15^N enrichment is presented as a percentage of the total metabolite, data show mean 95% confidence interval. Hatched bars indicate summer squirrels (n = 29), solid bars indicate hibernating squirrels (n= 6). Statistical analyses conducted only for metabolites detected in both hibernation and summer samples via multivariate linear regression, with diet as a covariate (p<0.05).

### ^15^N metabolites in cecal lumen contents and cecal tissue cluster separately from other tissues in hibernating and summer active squirrels

^15^N Metabolite profiles differed significantly between samples from hibernating and summer active squirrels, but only explained 2.3% of variation (PERMANOVA: F_1,201_=5.08, R^2^=0.023, p<0.001). Tissue type explained 87.7% of variation in ^15^N metabolite profiles in hibernating squirrels (PERMANOVA: F_5,33_=47.19, R^2^=0.8773, p<0.001), and 70.8% in summer active squirrels (PERMANOVA: F_5,167_=80.85, R^2^=0.7077, p<0.001) (Fig. 8). Diet did not significantly affect ^15^N metabolite profiles in either hibernating or summer active squirrels (p=0.957; p=0.639).

**Fig. 8.**
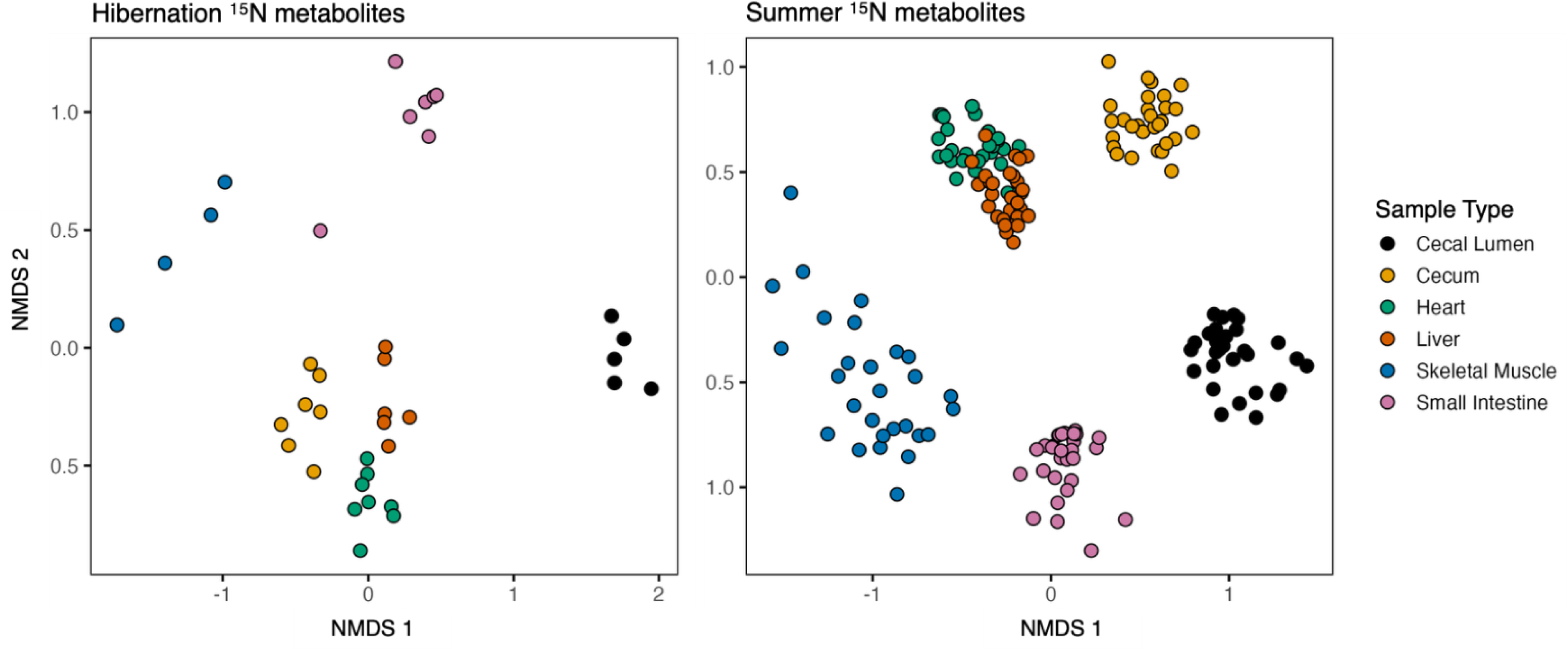
Non-metric Multidimensional Scaling (NMDS) using a Bray-Curtis dissimilarity distribution urea-derived ^15^N metabolite enrichment distinct clustering of heart and liver tissues in hibernating and summer active squirrels. Cecum tissues also clustered with heart and liver tissues during hibernation, but not in summer-active squirrels. Sample type explained 87.7% of variation in metabolite profiles in hibernating squirrels compared to 70.8% in summer-active squirrels (p<0.001).

## Discussion

We found ^15^N-enriched metabolites within the gastrointestinal tract and in peripheral tissues of both summer active and hibernating arctic ground squirrels, demonstrating incorporation of MLUN into both essential and nonessential AAs (including EBCAAs), as well as non-amino acid metabolites such as purines. Notably, we found substantially more MLUN incorporated into glutamine, citrulline, and leucine-isoleucine across multiple tissues in hibernating squirrels than in summer active squirrels, supporting our hypothesis that anabolic AA metabolism pathways utilizing MLUN are upregulated in hibernation (Fig. 9). The occurrence of numerous ^15^N labeled metabolites outside the microbially rich-GI tract indicates that host enzymatic mechanisms may facilitate the conversion of MLUN into critical nutrients.

**Fig. 9.**
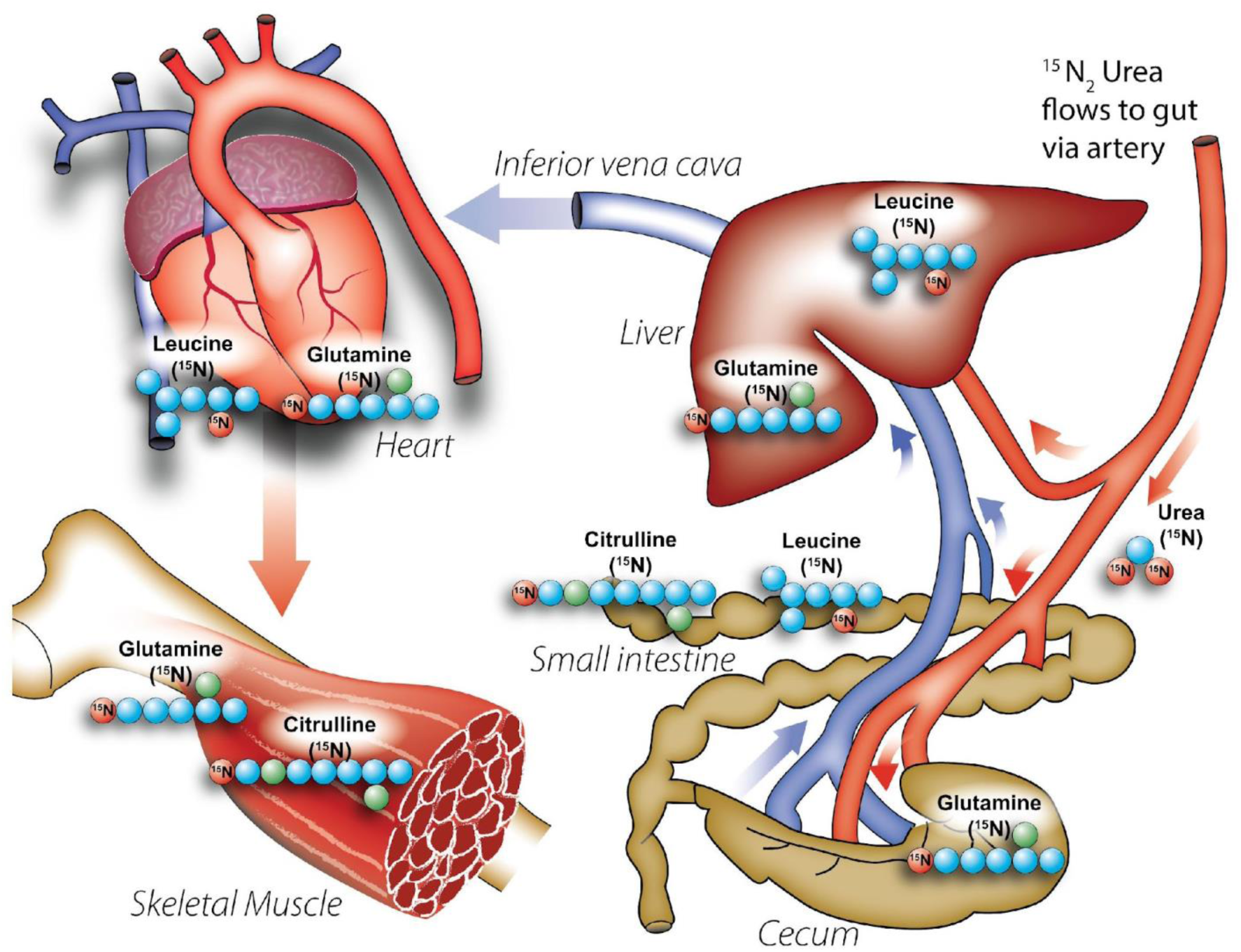
Microbially liberated ^15^N from urea appears across peripheral organs in ground squirrels. Microbially-derived urea nitrogen (^15^N) is increasingly incorporated into host tissues in hibernation and may be dependent on both the cecum as well as the small intestine

While we observed low concentrations (under 1%) of ^15^N-labeled threonine and methionine in heart and liver, indicating a possible link between cecal lumen microbial EAA *de novo* synthesis and uptake by host peripheral tissues, the level of ^15^N enrichment was not higher in hibernation. In fact, hepatic ^15^N methionine was higher in summer active squirrels, suggesting that this pathway is not upregulated in hibernation. In contrast to the lower EAA ^15^N incorporation, a high level of ^15^N enrichment of glutamine, citrulline, and leucine-isoleucine in hibernating squirrels (upwards of 10% in some gut and peripheral organs) is particularly striking given the unique role each amino acid plays in the GI tract, microbial, and whole-body nitrogen metabolism, as well as their strong influence on protein balance and insulin sensitivity ^30,55–57^.

Further, in both summer-active and hibernating squirrels, the metabolite profiles of heart and liver tissues clustered together. This finding could be attributed to the urea-nitrogen released by microorganisms in the cecum being rapidly absorbed into the portal bloodstream, and subsequently encountering the liver and then heart; these results could suggest that liberated ^15^N-ammonia is incorporated into metabolites via *de novo* synthesis pathways in liver and exported to the heart, leading to similar metabolite communities. During hibernation, cecum tissues also clustered with heart and liver tissues. This is likely due to the close proximity of the cecum tissue to the ^15^N generated by the cecal microbiome, and might indicate cecal and portal blood flow uptake of microbially-derived ^15^N-ammonia or ^15^N metabolites. Additionally, an increase in urea transporters in the cecum during hibernation (Regan et al. 2022) versus summer may contribute to this clustering and increased MLUN salvage^20^. Lastly, sample type explained more variation in metabolite profiles in hibernating squirrels than in summer-active squirrels. This increased variation during hibernation may be explained by the lack of dietary nitrogen intake, whereas in summer-active squirrels, dietary nitrogen intake could dilute the nitrogen signal, thereby reducing the observed variation in metabolite profiles.

### Urea-nitrogen recycling into NEAAs and EBCAAs is upregulated in hibernation

We propose the recycling of MLUN via leucine-isoleucine, glutamine, and citrulline synthesis may be favored specifically for their influence on protein balance, insulin sensitivity, and their role in nitrogen transport^30,55–57^. Such physiological roles are especially important in the ecological context of arctic ground squirrel hibernation; this species hibernates for long periods at sub-zero ambient temperatures during which they are thermogenic and fuel this process in part through tissue catabolism^12,13,33,46^. Due to the fact that arctic ground squirrels hibernating at lower ambient temperatures rely on amino acids as well as lipids to fuel metabolism^12^, mechanisms conserving nitrogen and maintaining protein balance may need to be increased. Multiple species show resistance to muscle loss^58,59^ and upregulate expression of genes related to anabolism^34^ in hibernation, and UNS has long been theorized to facilitate retention protein stores^14,18,20^. There is some alteration of AA and urea cycle gene expression when arctic ground squirrels hibernate at colder temperatures^60^, indicating AA metabolic pathways in hibernation may be responsive to thermal challenges; however, this has been a relatively unexplored field of study.

#### Citrulline

The small intestine is the primary site for systemic citrulline production^53^, and we found greater incorporation of MLUN into citrulline in the small intestine of hibernating vs. summer active squirrels. However, regardless of physiological state (hibernation vs summer active), the percent of ^15^N-labeled citrulline was an order of magnitude lower in the small intestine tissue and the cecal lumen contents as compared to cecal tissue, suggesting that epithelial cells of the cecum may be capable of citrulline production in arctic ground squirrels. In addition, there was a significantly higher percentage of ^15^N-citrulline in the tissues of the cecum and small intestine as compared to other NEAAs, suggesting that citrulline production may be an important initial product following liberation of ammonia from urea by the cecal microbiota. The appearance of ^15^N citrulline in the peripheral skeletal muscle surprised us, however, given that citrulline is not incorporated into structural tissue protein. ^15^N arginine, which is a major product of citrulline and stoichiometrically linked in the kidney^30^, also appeared in skeletal muscle and heart tissue (albeit in very modest amounts)^30^, and enzymes responsible for interconversion of both arginine and citrulline are present in endothelial cells across the body, including blood vessels^61,62^. Given this,^15^N arginine in peripheral skeletal muscle may represent a direct link between incorporation of MLUN into ^15^N citrulline in the GI tract, which then bypassed hepatic catabolism to be converted to arginine in the periphery.

Metabolic reactions are tightly regulated in hibernation^63,64^, so it is surprising that nitrogen recycling into a non-protein amino acid, such as citrulline, would be upregulated in hibernating arctic ground squirrels. Citrulline, however, is unique among amino acids because it can by-pass hepatic uptake (thereby avoiding hepatic catabolism) to reach peripheral tissues^53,65^, and as such has been proposed as an alternative carrier of nitrogen^56^. In rats, citrulline has been shown to 1) stimulate protein synthesis, 2) serve as a conditionally essential amino acid when intestinal function is compromised, and 3) restore nitrogen balance^56^. Curiously, we previously showed that hibernating squirrels provisioned with free ^15^N ammonium did not significantly increase ^15^N citrulline in most tissues^31^. This finding was supported by results of Regan et al. (2022) who also did not document this metabolite in their study, although it is unclear if they specifically tested for it^20,31^. Given the role of citrulline in maintaining nitrogen balance, and that arctic ground squirrels utilize mixed fuel metabolism and lose more lean mass when hibernating at ambient temperatures below zero ^12,33^, our contrasting results could be due to low hibernation temperatures of this study (−16°C), as compared to the thermal neutral hibernation temperatures used in our previous study and that of Regan et al^20,31^.

#### Leucine-isoleucine

We show that MLUN was incorporated into the EBCAAs leucine-isoleucine, with very high percentages in the liver, heart and small intestine of hibernating squirrels. We found no incorporation of MLUN into leucine-isoleucine in cecal tissue. Our results are consistent with the fact that a major role of EBCAAs in metabolism, in particular that of leucine, is to stimulate anabolism via the mammalian target of rapamycin (mTOR), a major signaling pathway for protein synthesis^30^. This also matches Lee et al.’s finding of protein anabolism in squirrel liver, heart and small intestine, but not skeletal muscle, during hibernation at sub-zero ambient temperatures^33^. Our evidence of *de novo* leucine-isoleucine production is also consistent with our previous work in hibernating squirrels^31^ showing that ^15^N from IV injected ^15^N ammonium was incorporated into leucine-isoleucine in multiple tissues, likely via BCAA aminotransferase activity. Our expanded time-scale experiments across natural torpor bouts using ^15^N_2_ urea in this study specifically highlights that BCAA transaminase activity may be increased in the liver, heart and small intestine of hibernating arctic ground squirrels.

The very high percentage of leucine-isoleucine in the liver as compared to any GI tract sample indicates that the majority of *de novo* synthesis of EBCAA may occur outside the GI tract via uptake of ^15^N ammonia (liberated from urea) and transaminase reactions; however, the gut microbiota may also be contributing directly to systemic host EBCAA pools. Gut microbes have high levels of EBCAAs in their protein^26,66^, and leucine synthesis from short-chain fatty acids has been documented in some gut bacteria^27^. Additionally, EBCAA biosynthesis and degradation pathways were more highly expressed in the metatranscriptome of the small intestine lumen microbiota than in the lumen microbiome of the cecum in the same hibernating individuals^45^. Our results of ^15^N leucine-isoleucine in small intestine, but not cecum, additionally support the supposition that microbial EBCAA biosynthesis most likely is occurring within the small intestine, not the cecum. Interestingly, Regan et al. documented in 13-lined ground squirrels that ^15^N incorporation into host proteins and metabolites during hibernation was lower in cecal tissue but higher in liver tissue as compared to during the summer active period^20^. In combination, our study and that of Regan et al. suggest that non-cecal tissue may contribute to UNS during hibernation in multiple species of hibernators, and future studies in a wider variety of hibernators are needed to support such a conclusion.

#### Glutamine

Given that glutamine is central to nitrogen transport systemically ^30^, we expected it to be a major contributor to inter-organ nitrogen salvage in fasting arctic ground squirrels during hibernation. Indeed, we found more MLUN incorporated into glutamine in the liver, heart, skeletal muscle and cecal tissue in hibernating animals than in active squirrels. Interestingly, we did not detect ^15^N glutamine in the small intestine of summer active squirrels; we found negligible amounts in the hibernators (Supplemental Data File 1), and we detected no ^15^N glutamine in any cecal lumen contents. Such results would indicate that the small intestine and cecal microbiota are not the source of endogenous glutamine production. Additionally, in the cecal lumen content we saw significant enrichment of MLUN in glutamate and other metabolites such as 5-oxoproline, with glutamate being higher in hibernating than in summer-active squirrels. Therefore, evidence indicates that glutamine production for systemic circulation is dependent on either the cecal enterocytes which derive nitrogen from either free ammonia liberated by urease or microbially generated ^15^N metabolites (e.g.,^15^N-glutamate) produced in the cecal lumen, or from glutamine production via MLUN occurring outside the GI tract.

#### Purine and pyrimidine nitrogen recycling and other metabolites

The story of nitrogen salvage in hibernation often revolves around AAs, but purines and pyrimidines are major nitrogen carrying metabolites as well. In other species, such as elephant seals, fasting increases purine recycling^67^, and we have shown previously that purine end-products accumulate as a torpor bout progresses ^31^. Hibernation biochemistry and physiology is fundamentally geared toward conserving energy and resources and our observances of MLUN into purine and pyrimidine nucleoside and nucleobase metabolism may indicate an under-studied mechanism to support cellular function in long-term fasting.

In addition to purines and pyrimidines, ^15^N was incorporated into other metabolites, such as alanine, lysine, phenylalanine, tryptophan, spermidine, 5-oxoproline, glycine and serine. ^15^N alanine was higher in hibernating animals across the GI tract and liver. Given that alanine is the major amino acid contributing to gluconeogenesis in liver^68^ and arctic ground squirrels engage in gluconeogenesis during arousal from torpor^7^, it is intriguing to consider whether alanine synthesis is preferentially upregulated during thermogenic torpor to supply a gluconeogenic substrate.

### Additional considerations for nitrogen recycling in hibernators

We were surprised by the degree to which MLUN was incorporated into tissue in hibernating vs summer active squirrels, especially considering that both anabolic and catabolic processes are highly depressed in ground squirrels during hibernation ^12,69^. In our study, hibernating arctic ground squirrels were maintained at an ambient temperature of −16°C, an ambient temperature that requires increase of torpid-MR from 1% of basal metabolic rate to a thermogenic output of ∼50% of basal metabolic rate in order to maintain body temperature >−2.9°C^12,38,46^. While the relatively high metabolic rate required to maintain body temperature at a low ambient temperature of hibernation may help explain this discrepancy, the percent enrichment of isotopically labeled urea to the existing urea pools in summer active vs hibernating squirrels must be taken into consideration when interpreting our results.

In obligate seasonal hibernators, such as ground squirrels, the turnover rates of total body water (TBW) are lower during hibernation than during summer euthermia because animals neither eat nor drink during hibernation, do not urinate while torpid (as kidney function is highly depressed^70,71^),excrete little urine during IBAs^72,73^, and reduce water loss via lower metabolic rates in torpor^74^. Consequently, more isotopically labeled urea, which rapidly equilibrates with TBW^75^, might be lost in summer active compared to hibernating animals, altering the percent enrichment between groups. Additionally, plasma urea concentrations in squirrels^8^ and bears^76^ are lower in hibernation than summer euthermia. The majority of isotope tracer studies conducted in hibernating animals have measured levels of isotopic incorporation rather than kinetics, and used the same dose of isotope for both summer active and hibernating animals ^20,77^. Given the overall lower urea concentrations in hibernation, injected isotopically labeled urea may enrich a larger percentage of overall urea pool in hibernators compared to summer animals, enhancing overall percentage of ^15^N recycling from urea in hibernation. While both of these facets are important in comparing summer to hibernation urea recycling in hibernation and should be clarified with future studies, it is important to note how depressed all metabolic processes are in hibernation and how astonishing it is that metabolic recycling of urea nitrogen into different metabolic pathways would be upregulated across the body. Additionally, it is important to note, reduction of urea loss to low TBW loss in itself is a potential nitrogen sparing mechanism, naturally giving microbiota more time to salvage urea nitrogen, as studies show decreased urinary water loss and increased urea degradation in the gut during periods of water restriction in ruminants and hindgut fermenters ^16,78,79^.

## Conclusion

Our results indicate a possible influence of microbially-liberated urea-nitrogen supporting nitrogen recycling in multiple host organs, and upregulated *de novo* citrulline, glutamine and leucine-isoleucine synthesis in hibernation. We therefore propose that upregulated UNS into synthesis of anabolic amino acids, such as EBCAAs and citrulline, may play an important role in conserving protein balance in hibernation, and may influence hibernator physiology and metabolism, especially when hibernating at thermogenic ambient temperatures. Further, our results highlight that the small intestine may play a significant role in contributing to host uptake of MLUN associated metabolites and deserves further investigation.

A remaining question, however, is whether these metabolites derive from ^15^N incorporation into microbial AAs and protein, or from direct ammonia release from microbes to the cecum or small intestine epithelium. Future studies using *in vivo* techniques to directly sample from the portal and hepatic veins of hibernators would greatly enrich our understanding of how MLUN is involved in host metabolism and potentially help answer such questions. Such cannulation techniques have been used in pigs, mice, and humans ^80–82^ and could be employed in hibernators. Additionally, future studies quantifying ecologically relevant rates of protein synthesis in tissues of hibernators would contribute to understanding and quantifying proportions of MLUN metabolites that are incorporated into protein and the impact of MLUN on protein pools, as there have been few *in vivo* protein synthesis studies in hibernators ^83,84^.

## Acknowledgments

We would like to thank Karen Carlson, Erica Chenoweth, Bailey Fuller, Jasmine Hatton, Jordan Houk, Sherri Hughes, Alyssa Igtanloc, Travis Jennings, Emma Luck, Shannon Medlock, Ashley Mizell, Kat O’Brien, Joyce Pexton, Emily Shibe, Ashlee Voorhis, and Frank Woitel for assistance with animal husbandry and experiments, and Francesca Cendali and Rebecca Wilkerson for assistance with metabolomics data acquisition.

## Author Contributions

Conceived and designed the experiments: KD, CLB

Performed the experiments: SG, KG

Analyzed the data: SR, JH, KD

Contributed materials/analysis tools: KD, TB-P, JH, SR

Wrote the paper: SR, KD

All authors contributed to editing the manuscript

## Abbreviations

IP: intraperitoneal
AA: amino acid
EAA: essential amino acid
NEAA: non-essential amino acid
EBCAA: essential branched-chain amino acid
UNS: urea nitrogen salvage
MLUN: microbially-liberated urea-nitrogen
GI tract: gastrointestinal tract
IBA: interbout arousal

## References

1 Turbill, C., Bieber, C. & Ruf, T. Hibernation is associated with increased survival and the evolution of slow life histories among mammals. Proc Biol Sci 278, 3355–3363, doi:10.1098/rspb.2011.0190 (2011).

2 Carey, H. V., Andrews, M. T. & Martin, S. L. Mammalian hibernation: cellular and molecular responses to depressed metabolism and low temperature. Physiol Rev 83, 1153–1181, doi:10.1152/physrev.00008.2003 (2003).

3 Heldmaier, G., Ortmann, S. & Elvert, R. Natural hypometabolism during hibernation and daily torpor in mammals. Respir Physiol Neurobiol 141, 317–329, doi:10.1016/j.resp.2004.03.014 (2004).

4 Boyer, B. B. & Barnes, B. M. Molecular and Metabolic Aspects of Mammalian Hibernation. BioScience 49, 713–724 (1999).

5 Barnes, B. M. & Ritter, D. in Life in the Cold: Ecological, Physiological, and Molecular Mechanisms (ed C.; Florant Carey, G.L., Wunder, B.A.; Horwitz, B.) (Westview Press, 1993).

6 van Breukelen, F. & Martin, S. L. Translational initiation is uncoupled from elongation at 18 degrees C during mammalian hibernation. Am J Physiol Regul Integr Comp Physiol 281, R1374–1379, doi:10.1152/ajpregu.2001.281.5.R1374 (2001).

7 Galster, W. & Morrison, P. R. Gluconeogenesis in arctic ground squirrels between periods of hibernation. Am J Physiol 228, 325–330, doi:10.1152/ajplegacy.1975.228.1.325 (1975).

8 Epperson, L. E., Karimpour-Fard, A., Hunter, L. E. & Martin, S. L. Metabolic cycles in a circannual hibernator. Physiol Genomics 43, 799–807, doi:10.1152/physiolgenomics.00028.2011 (2011).

9 Barnes, B. M. Freeze avoidance in a mammal: body temperatures below 0 degree C in an Arctic hibernator. Science 244, 1593–1595 (1989).

10 Drew, K. L. et al. Central nervous system regulation of mammalian hibernation: implications for metabolic suppression and ischemia tolerance. J Neurochem 102, 1713–1726, doi:10.1111/j.1471-4159.2007.04675.x (2007).

11 Bogren, L. K. et al. The effects of hibernation and forced disuse (neurectomy) on bone properties in arctic ground squirrels. Physiol Rep 4, doi:10.14814/phy2.12771 (2016).

12 Buck, C. L. & Barnes, B. M. Effects of ambient temperature on metabolic rate, respiratory quotient, and torpor in an arctic hibernator. Am J Physiol Regul Integr Comp Physiol 279, R255–262, doi:10.1152/ajpregu.2000.279.1.R255 (2000).

13 Buck, C. L. B., B.M. Temperatures of Hibernacula and Changes in Body Composition of Arctic Ground Squirrels over Winter. Journal of Mammology 80, 1264–1276 (1999).

14 Barboza, P. S., Farley, S. D. & Robbins, C. T. Whole-body urea cycling and protein turnover during hyperphagia and dormancy in growing bears (Ursus americanus and U. arctos). Canadian Journal of Zoology 75, 2129–2136, doi:10.1139/z97-848 (1997).

15 Stewart, G. S. & Smith, C. P. Urea nitrogen salvage mechanisms and their relevance to ruminants, non-ruminants and man. Nutr Res Rev 18, 49–62, doi:10.1079/NRR200498 (2005).

16 Stevens, C. E. & Hume, I. D. Contributions of microbes in vertebrate gastrointestinal tract to production and conservation of nutrients. Physiol Rev 78, 393–427, doi:10.1152/physrev.1998.78.2.393 (1998).

17 Mobley, H. L. & Hausinger, R. P. Microbial ureases: significance, regulation, and molecular characterization. Microbiol Rev 53, 85–108, doi:10.1128/mr.53.1.85-108.1989 (1989).

18 Steffen, J. M., Rigler, G. L., Moore, A. K. & Riedesel, M. L. Urea recycling in active golden-mantled ground squirrels (Spermophilus lateralis). Am J Physiol 239, R168–173, doi:10.1152/ajpregu.1980.239.1.R168 (1980).

19 Nelson, R. A. Urea metabolism in the hibernating black bear. Kidney Int Suppl, S177–179 (1978).

20 Regan, M. D. et al. Nitrogen recycling via gut symbionts increases in ground squirrels over the hibernation season. Science 375, 460–463, doi:10.1126/science.abh2950 (2022).

21 Sadowska, J., Carlson, K. M., Buck, C. L., Lee, T. N. & Duddleston, K. N. Microbial urea-nitrogen recycling in arctic ground squirrels: the effect of ambient temperature of hibernation. J Comp Physiol B 194, 909–924, doi:10.1007/s00360-024-01579-9 (2024).

22 Millward, D. J. et al. The transfer of 15N from urea to lysine in the human infant. Br J Nutr 83, 505–512 (2000).

23 Metges, C. C. et al. Incorporation of urea and ammonia nitrogen into ileal and fecal microbial proteins and plasma free amino acids in normal men and ileostomates. Am J Clin Nutr 70, 1046–1058, doi:10.1093/ajcn/70.6.1046 (1999).

24 Regoeczi, E., Irons, L., Koj, A. & McFarlane, A. S. Isotopic Studies of Urea Metabolism in Rabbits. Biochem J 95, 521–532, doi:10.1042/bj0950521 (1965).

25 Davila, A. M. et al. Intestinal luminal nitrogen metabolism: role of the gut microbiota and consequences for the host. Pharmacol Res 68, 95–107, doi:10.1016/j.phrs.2012.11.005 (2013).

26 Neis, E. P., Dejong, C. H. & Rensen, S. S. The role of microbial amino acid metabolism in host metabolism. Nutrients 7, 2930–2946, doi:10.3390/nu7042930 (2015).

27 Portune, K. J. B., M; Davila, A.; Tome, D.; Blachier, F.; Sanz, Y. Gut microbiota role in dietary protein metabolism and health-related outcomes: The two sides of the coin. Trends Food Sci. Technol. 57, 213–232 (2016).

28 van de Poll, M. C. et al. Intestinal and hepatic metabolism of glutamine and citrulline in humans. J Physiol 581, 819–827, doi:10.1113/jphysiol.2006.126029 (2007).

29 van de Poll, M. C. et al. Interorgan amino acid exchange in humans: consequences for arginine and citrulline metabolism. Am J Clin Nutr 85, 167–172, doi:10.1093/ajcn/85.1.167 (2007).

30 Cynober, L. A. Metabolic and Therapeutic Aspects of Amino Acids in Clinical Nutrition. 2nd edn, (CRC Press, 2003).

31 Rice, S. A. et al. Nitrogen recycling buffers against ammonia toxicity from skeletal muscle breakdown in hibernating arctic ground squirrels. Nat Metab 2, 1459–1471, doi:10.1038/s42255-020-00312-4 (2020).

32 Chanon, S. et al. Proteolysis inhibition by hibernating bear serum leads to increased protein content in human muscle cells. Sci Rep 8, 5525, doi:10.1038/s41598-018-23891-5 (2018).

33 Lee, T. N., Buck, C. L., Barnes, B. M. & O’Brien, D. M. A test of alternative models for increased tissue nitrogen isotope ratios during fasting in hibernating arctic ground squirrels. J Exp Biol 215, 3354–3361, doi:10.1242/jeb.068528 (2012).

34 Fedorov, V. B. et al. Comparative functional genomics of adaptation to muscular disuse in hibernating mammals. Mol Ecol 23, 5524–5537, doi:10.1111/mec.12963 (2014).

35 Fedorov, V. B. et al. Modulation of gene expression in heart and liver of hibernating black bears (Ursus americanus). BMC Genomics 12, 171, doi:10.1186/1471-2164-12-171 (2011).

36 Bertile, F., Habold, C., Le Maho, Y. & Giroud, S. Body Protein Sparing in Hibernators: A Source for Biomedical Innovation. Front Physiol 12, 634953, doi:10.3389/fphys.2021.634953 (2021).

37 Wolfe, R. R., Nelson, R. A., Wolfe, M. H. & Rogers, L. Nitrogen cycling in hibernating bears. Proc. Amer. Soc. Mass Spectrometry 426 (1982).

38 Richter, M. M. et al. Thermogenic capacity at subzero temperatures: how low can a hibernator go? Physiol Biochem Zool 88, 81–89, doi:10.1086/679591 (2015).

39 Buck, C. L., Breton, A., Kohl, F., Toien, O. & Barnes, B. M. in Hypometabolism in animals: torpor hibernation and cryobiology 317–326 (2008).

40 Doolin, M. L. & Dearing, M. D. Differential Effects of Two Common Antiparasitics on Microbiota Resilience. J Infect Dis 229, 908–917, doi:10.1093/infdis/jiad547 (2024).

41 Carey, H. V., Walters, W. A. & Knight, R. Seasonal restructuring of the ground squirrel gut microbiota over the annual hibernation cycle. Am J Physiol Regul Integr Comp Physiol 304, R33–42, doi:10.1152/ajpregu.00387.2012 (2013).

42 Stevenson, T. J., Duddleston, K. N. & Buck, C. L. Effects of season and host physiological state on the diversity, density, and activity of the arctic ground squirrel cecal microbiota. Appl Environ Microbiol 80, 5611–5622, doi:10.1128/AEM.01537-14 (2014).

43 Stevenson, T. J., Buck, C. L. & Duddleston, K. N. Temporal dynamics of the cecal gut microbiota of juvenile arctic ground squirrels: a strong litter effect across the first active season. Appl Environ Microbiol 80, 4260–4268, doi:10.1128/AEM.00737-14 (2014).

44 Pengelley, E. T. & Fisher, K. C. Rhythmical arousal from hibernation in the golden-mantled ground squirrel, *Citellus lateralis* tescorum. Canadian Journal of Zoology 39, 105–120, doi:10.1139/z61-013 (1961).

45 Grond, K., Buck, C. L. & Duddleston, K. N. Microbial gene expression during hibernation in arctic ground squirrels: greater differences across gut sections than in response to pre-hibernation dietary protein content. Front Genet 14, 1210143, doi:10.3389/fgene.2023.1210143 (2023).

46 Karpovich, S. A., Toien, O., Buck, C. L. & Barnes, B. M. Energetics of arousal episodes in hibernating arctic ground squirrels. J Comp Physiol B 179, 691–700, doi:10.1007/s00360-009-0350-8 (2009).

47 Nemkov, T., Hansen, K. C. & D’Alessandro, A. A three-minute method for high-throughput quantitative metabolomics and quantitative tracing experiments of central carbon and nitrogen pathways. Rapid Commun Mass Spectrom 31, 663–673, doi:10.1002/rcm.7834 (2017).

48 Gehrke, S. et al. Red Blood Cell Metabolic Responses to Torpor and Arousal in the Hibernator Arctic Ground Squirrel. J Proteome Res, doi:10.1021/acs.jproteome.9b00018 (2019).

49 Nemkov, T., Reisz, J. A., Gehrke, S., Hansen, K. C. & D’Alessandro, A. High-Throughput Metabolomics: Isocratic and Gradient Mass Spectrometry-Based Methods. Methods Mol Biol 1978, 13–26, doi:10.1007/978-1-4939-9236-2_2 (2019).

50 Kotera, M., Hirakawa, M., Tokimatsu, T., Goto, S. & Kanehisa, M. The KEGG databases and tools facilitating omics analysis: latest developments involving human diseases and pharmaceuticals. Methods Mol Biol 802, 19–39, doi:10.1007/978-1-61779-400-1_2 (2012).

51 Kanehisa, M. The KEGG database. Novartis Found Symp 247, 91–101; discussion 101-103, 119-128, 244-152 (2002).

52 Xia, J. & Wishart, D. S. Web-based inference of biological patterns, functions and pathways from metabolomic data using MetaboAnalyst. Nat Protoc 6, 743–760, doi:10.1038/nprot.2011.319 (2011).

53 Windmueller, H. G. & Spaeth, A. E. Source and fate of circulating citrulline. Am J Physiol 241, E473–480, doi:10.1152/ajpendo.1981.241.6.E473 (1981).

54 Trommelen, J. & van Loon, L. J. C. Assessing the whole-body protein synthetic response to feeding in vivo in human subjects. Proc Nutr Soc 80, 139–147, doi:10.1017/S0029665120008009 (2021).

55 Rennie, M. J. et al. Characteristics of a glutamine carrier in skeletal muscle have important consequences for nitrogen loss in injury, infection, and chronic disease. Lancet 2, 1008–1012, doi:10.1016/s0140-6736(86)92617-6 (1986).

56 Moinard, C. & Cynober, L. Citrulline: a new player in the control of nitrogen homeostasis. J Nutr 137, 1621S–1625S, doi:10.1093/jn/137.6.1621S (2007).

57 Ham, D. J. et al. L-Citrulline Protects Skeletal Muscle Cells from Cachectic Stimuli through an iNOS-Dependent Mechanism. PLoS One 10, e0141572, doi:10.1371/journal.pone.0141572 (2015).

58 Cotton, C. J. & Harlow, H. J. Avoidance of skeletal muscle atrophy in spontaneous and facultative hibernators. Physiol Biochem Zool 83, 551–560, doi:10.1086/650471 (2010).

59 Lohuis, T. D., Harlow, H. J., Beck, T. D. & Iaizzo, P. A. Hibernating bears conserve muscle strength and maintain fatigue resistance. Physiol Biochem Zool 80, 257–269, doi:10.1086/513190 (2007).

60 Williams, C. T. et al. Hibernating above the permafrost: effects of ambient temperature and season on expression of metabolic genes in liver and brown adipose tissue of arctic ground squirrels. J Exp Biol 214, 1300–1306, doi:10.1242/jeb.052159 (2011).

61 Husson, A., Brasse-Lagnel, C., Fairand, A., Renouf, S. & Lavoinne, A. Argininosuccinate synthetase from the urea cycle to the citrulline-NO cycle. Eur. J. Biochem. 270, 1887–1899, doi:10.1046/j.1432-1033.2003.03559.x (2003).

62 Wu, G. & Morris, S. M., Jr. Arginine metabolism: nitric oxide and beyond. Biochem J 336 (Pt 1), 1–17 (1998).

63 Storey, K. B. & Storey, J. M. Metabolic rate depression and biochemical adaptation in anaerobiosis, hibernation and estivation. Q Rev Biol 65, 145–174, doi:10.1086/416717 (1990).

64 Storey, K. B. Metabolic regulation in mammalian hibernation: enzyme and protein adaptations. Comp Biochem Physiol A Physiol 118, 1115–1124, doi:10.1016/s0300-9629(97)00238-7 (1997).

65 Bahri, S. et al. Citrulline: from metabolism to therapeutic use. Nutrition 29, 479–484, doi:10.1016/j.nut.2012.07.002 (2013).

66 Diether, N. E. & Willing, B. P. Microbial Fermentation of Dietary Protein: An Important Factor in Diet(-)Microbe(-)Host Interaction. Microorganisms 7, doi:10.3390/microorganisms7010019 (2019).

67 Sonanez-Organis, J. G. et al. Prolonged fasting increases purine recycling in post-weaned northern elephant seals. J Exp Biol 215, 1448–1455, doi:10.1242/jeb.067173 (2012).

68 Stumvoll, M. et al. Human kidney and liver gluconeogenesis: evidence for organ substrate selectivity. Am J Physiol 274, E817–826, doi:10.1152/ajpendo.1998.274.5.E817 (1998).

69 Geiser, F. Metabolic rate and body temperature reduction during hibernation and daily torpor. Annu Rev Physiol 66, 239–274, doi:10.1146/annurev.physiol.66.032102.115105 (2004).

70 Moy, R. M. Renal function in the hibernating ground squirrel Spermophilus columbianus. Am J Physiol 220, 747–753, doi:10.1152/ajplegacy.1971.220.3.747 (1971).

71 Lesser, R. W., Moy, R., Passmore, J. C. & Pfeiffer, E. W. Renal regulation of urea excretion in arousing and homeothermic ground squirrels (Citellus columbianus). Comp Biochem Physiol 36, 291–296, doi:10.1016/0010-406x(70)90009-5 (1970).

72 Passmore, J. C., Pfeiffer, E. W. & Templeton, J. R. Urea excretion in the hibernating Columbian ground squirrel (Spermophilus columbianus). J Exp Zool 192, 83–86, doi:10.1002/jez.1401920110 (1975).

73 Pengelley, E. T., Asmundson, S. J., Aloia, R. C. & Barnes, B. Circannual rhythmicity in a non-hibernating ground squirrel, Citellus leucurus. Comp Biochem Physiol A Comp Physiol 54, 233–237, doi:10.1016/s0300-9629(76)80103-x (1976).

74 Fisher, K. & Mannery, J. (eds KC; Fisher, AP; Dawe, & CP Lyman) 235–279 (Elsevier, Mammalian Hibernation III, 1967).

75 Matthews, D. E. & Downey, R. S. Measurement of urea kinetics in humans: a validation of stable isotope tracer methods. Am J Physiol 246, E519–527, doi:10.1152/ajpendo.1984.246.6.E519 (1984).

76 Stenvinkel, P. et al. Metabolic changes in summer active and anuric hibernating free-ranging brown bears (Ursus arctos). PLoS One 8, e72934, doi:10.1371/journal.pone.0072934 (2013).

77 Nelson, R. A. & Jones, J. D. Leucine Metabolism in the Black Bear. A Selection of Papers from the Seventh International Conference on Bear Research and Management, 329–331 (1986).

78 Freudenberger, D. O. & Hume, I. D. Effects of water restriction on digestive function in two macropodid marsupials from divergent habitats and the feral goat. J Comp Physiol B 163, 247–257, doi:10.1007/BF00261672 (1993).

79 Hume, I. D. R., K.; Engelhardt, W.V. Nitrogen Metabolism and Urea Kinetics in the Rock Hyrax (*Procavia habessinica*). Journal of Comparative Physiology B 138, 307–314 (1980).

80 Ten Have, G. A., Bost, M. C., Suyk-Wierts, J. C., van den Bogaard, A. E. & Deutz, N. E. Simultaneous measurement of metabolic flux in portally-drained viscera, liver, spleen, kidney and hindquarter in the conscious pig. Lab Anim 30, 347–358, doi:10.1258/002367796780739862 (1996).

81 Mercer, K. E. et al. Net release and uptake of xenometabolites across intestinal, hepatic, muscle, and renal tissue beds in healthy conscious pigs. Am J Physiol Gastrointest Liver Physiol 319, G133–G141, doi:10.1152/ajpgi.00153.2020 (2020).

82 Hallemeesch, M. M., Ten Have, G. A. & Deutz, N. E. Metabolic flux measurements across portal drained viscera, liver, kidney and hindquarter in mice. Lab Anim 35, 101–110, doi:10.1258/0023677011911426 (2001).

83 Whitten, B. K. & Klain, G. J. Protein metabolism in hepatic tissue of hibernating and arousing ground squirrels. Am J Physiol 214, 1360–1362, doi:10.1152/ajplegacy.1968.214.6.1360 (1968).

84 Hindle, A. G. et al. Prioritization of skeletal muscle growth for emergence from hibernation. J Exp Biol 218, 276–284, doi:10.1242/jeb.109512 (2015).

