## Supplemental Table and Figure for "Incorporation of microbially salvaged urea-nitrogen into anabolic amino acids during hibernation in arctic ground squirrels"

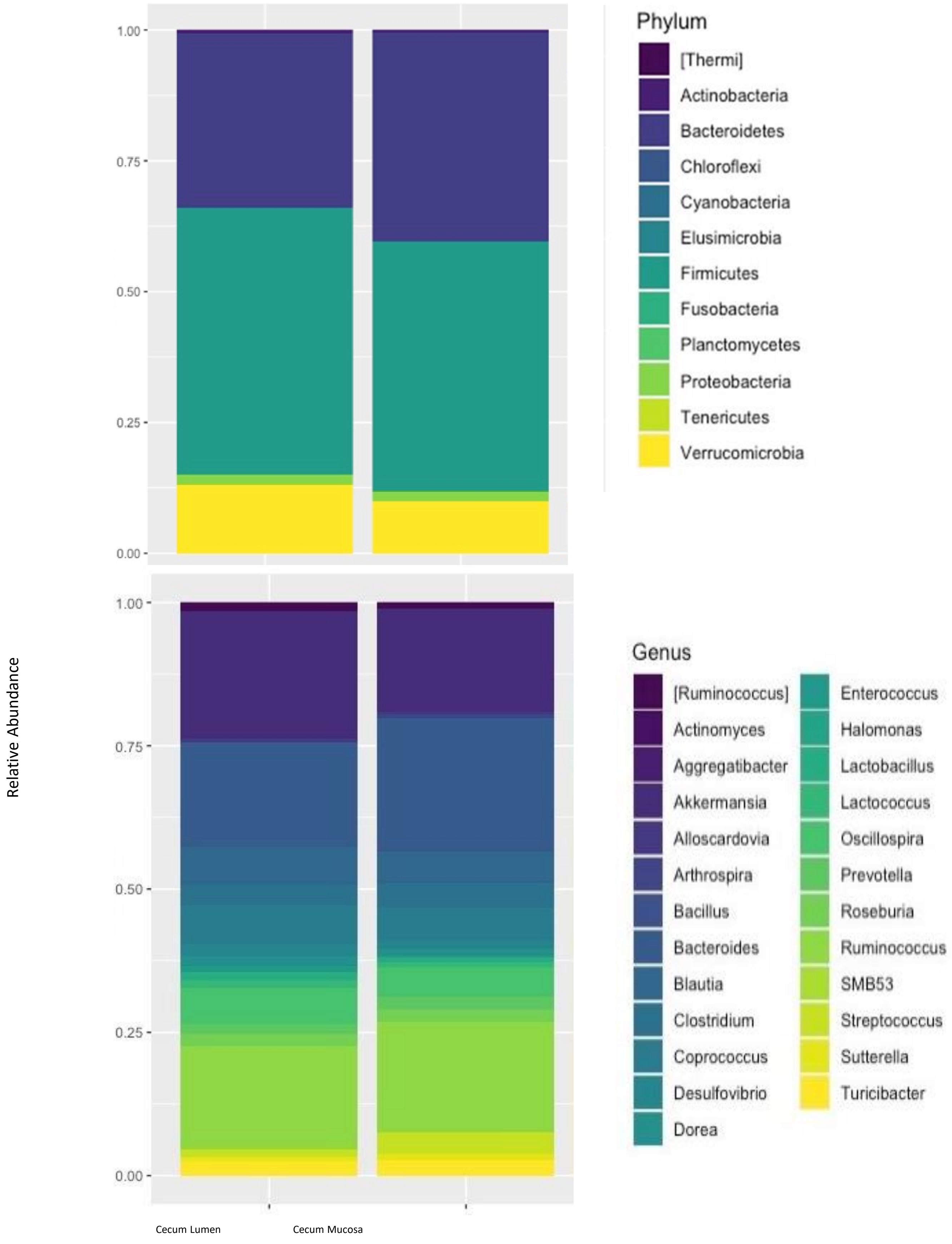


**Supplemental Figure 1.** Relative abundance of dominant taxa in the cecal microbial communities of summer active arctic ground squirrels. Top panel: Phylum. Bottom panel: Genus (top 25 most abundant genera). Diet and sex groups are combined within each gut compartment.

**Supplemental Table 1:** Effect of the addition of Karo syrup or sunflower oil on Kcal and macronutrient content of the diets

**2017-trapped squirrels**

50g pellets + 0.5g Karo syrup/day

9% protein diet

| | %Kcal from | | % by weight | |
|---|---|---|---|---|
| | no Karo | Karo | no Karo | Karo |
| Protein | 9.6 | 9.5 | 8.9 | 8.8 |
| Carb | 73.2 | 73.5 | 67.9 | 68.2 |
| Fat | 17.2 | 17 | 7.1 | 7.03 |

**2017-trapped squirrels**

50g pellets + 0.5g Karo syrup/day

18% protein diet

| | %Kcal from | | % by weight | |
|---|---|---|---|---|
| | no Karo | Karo | no Karo | Karo |
| Protein | 19.2 | 19 | 17.8 | 17.6 |
| Carb | 63.3 | 63.7 | 58.7 | 59.1 |
| Fat | 17.5 | 17.3 | 7.2 | 7.1 |

**2019-trapped squirrels**

45g pellets + 0.92g sunflower oil/day

9% protein diet

| | %Kcal from | | % by weight | |
|---|---|---|---|---|
| | no oil | oil | no oil | oil |
| Protein | 9.6 | 9.1 | 8.9 | 8.7 |
| Carb | 73.2 | 69.73 | 67.9 | 66.6 |
| Fat | 17.2 | 21.1 | 7.1 | 8.9 |

**2019-trapped squirrels**

45g pellets + 0.92g sunflower oil/day

18% protein diet

| | % Kcal from | | % by weight | |
|---|---|---|---|---|
| | no oil | oil | no oil | oil |
| Protein | 19.2 | 18.2 | 17.8 | 17.4 |
| Carb | 63.3 | 60.3 | 58.7 | 57.5 |
| Fat | 17.5 | 21.4 | 7.2 | 9.06 |
